# Quinolone resistance in different uropathogens isolated from urine cultures in patients with urinary tract infection

**DOI:** 10.1101/2020.09.07.286674

**Authors:** Jorge Angel Almeida Villegas, Iris Mellolzy Estrada Carrillo, Rodolfo Garcia Contreras, Silvia Patricia Peña

## Abstract

**Objective:** To identify patterns of resistance against quinolones in various uropathogens in urinary tract infections in the population of the Toluca valley, Mexico

**Introduction:** Quinolones are antibiotics with a spectrum of activity for both gram-positives and gram-negatives and are antibiotics used for the empirical treatment of urinary tract infections. Recently, a high index of resistance to quinolones has been reported due to different mechanisms on the part of bacteria, however the one that has taken the greatest importance is the presence of extended spectrum beta-lactamases

**Methods:** 155 samples were collected from patients with suspected urinary tract infection without exclusion criteria such as age or gender. Automated equipment was used for the identification of the etiological agent and sensitivity tests to quinolones.

**Results:** The results positives were divided to evaluate which of the two antibiotics studied had greater resistance. For ciprofloxacin there are 27 resistant strains 37%, 1 strain with intermediate resistance 1% and 45 susceptible strains 62%. For levofloxacin 26 strains are resistant 36%, 41 strains are sensitive 56% and 6 strains show intermediate sensitivity 8%.

**Conclusion:** Different microorganisms, both gram-positive and gram-negative, were isolated and it can be observed that gram-negative strains are the ones with the greatest resistance against quinolones, mainly *Escherichia coli*, which produces extended-spectrum beta-lactamases, in the case of gram-positive resistance patterns are variable with a tendency towards sensitivity.

## Introduction

Urinary tract infection (UTI) is the nosocomial infection that occurs most frequently worldwide^1^, being diagnosed both in outpatients and in hospitalized patients^2^. Affecting people of all ages, from newborn to geriatric age group^3^. Often, UTI patients have recurrent infections or failed empirical outpatient treatment, underlying comorbidities, or infections associated with medical care ^4^ The severity of the disease and the outcome of the UTI differ significantly and are related to several factors, including the sex, age, genetics and susceptibility of the host, the type of causative organism, the response to antibiotic therapy, the pattern of resistance to antibiotics and the clinical management of the patient. For this reason, urinalysis, culture, and antimicrobial susceptibility tests should be urgently performed on urine samples, as all this can be helpful in the diagnosis of UTI^5^. Among the most common causative agents identified are *Escherichia coli*, followed by *Klebsiella pneumoniae, Enterococcus faecalis, Proteus mirabilis, Pseudomonas aeruginosa, group B streptococci, Staphylococcus saprophyticus, Staphylococcus aureus*, and *Candida spp.*^6^ ^7^ ^8^ ^9^.

Currently, quinolones, which are a group of antibiotics classified into four generations with expanded activity for Gram-negative organisms as Gram-positive organisms^10^, are the third most prescribed antimicrobial drugs, followed by macrolides and beta-lactams for the treatment of various pathogenic infections^11^. Most cases of community-acquired UTI are treated empirically with quinolones without knowing the isolate’s pattern of drug susceptibility. Especially the use of Fluoroquinolones such as levofloxacin or ciprofloxacin, for complicated cases or UTI associated with catheters. Their increasing use in recent years has facilitated the emergence of quinolone-resistant uropathogens, which makes the treatment of infections caused by extended-spectrum β-lactamase-producing Enterobacteriaceae (ESBL) very limited^12^. Since ESBL genes are often encoded in plasmids, where they can also carry genes that encode resistance to other classes of antimicrobials, including quinolones^13^. Thus, it has been reported that resistance to quinolones is increasing, although these resistance rates show significant geographical variations^14^ ^15^.

Due to the rapid adaptation of bacteria, antimicrobial treatment for the management of urinary tract infections is becoming increasingly difficult, causing the antimicrobial profile of bacterial uropathogens to change over the years, despite having better antimicrobial agents. Resistance to these has increased significantly due to their incorrect use, limiting therapeutic options.^16^ Being a public health problem, especially in developing countries like Mexico, where, in addition to the high level of poverty, ignorance and poor hygiene practices, it is one of the concerns of medical society because treatment is expensive and often fails.^17^ For all of this, physicians must know the etiology and susceptibility patterns of urinary pathogens in their community, in order to adequately improve antimicrobial therapy ^18^ ^19^. Given that, bacterial pathogens and their patterns of susceptibility to antibiotics show particular differences by regions, finding that regional studies are much more valuable than international data^20^. To also carry out a timely diagnosis and therapy, preventing acute conditions that pose a risk to life, such as urosepsis, abscesses and permanent kidney scarring, which in turn could result in poor kidney growth, reduced glomerular function, hypertension arterial and chronic kidney disease^21^.

Consequently, local and current knowledge of uropathogens and their susceptibility and resistance patterns are of vital importance for an adequate selection and use of antibiotics, as well as for an adequate prescription policy. Therefore, the objective of this study was to determine the local antimicrobial resistance and susceptibility patterns to quinolones of uropathogens that were isolated in the Toluca, Mexico.

## Methods

In the present study, 155 urine cultures were performed in patients with clinical suspicion of urinary tract infection, without exclusion criteria such as gender or age. In a town in the center of the Toluca Valley, Mexico.

Of the 155 urine cultures performed, the distribution was; 107 people of the female gender representing 69% of the study population and 48 men representing 31%. The age ranges in the lower limit is a 1-year-old female patient and for the higher limit is a 91-year-old patient, with an average of 30 years.

For the study, 100 ml of the first urine of the day were collected with the necessary hygiene conditions for the collection of the specimen.

An aliquot was taken with a calibrated handle, and blood agar was inoculated to perform the colony count and chromogenic agar. Both Petri dishes were incubated for 24 hours at 37°C.

A colony isolated from the chromogenic agar medium was taken and resuspended in diluent medium, which was placed on the plates of the automated equipment.

The identification of the etiological agent as well as the sensitivity tests were performed using an automated method using the walkaway SI 96 from beckman coulter equipment.

## Results

Of the urine cultures obtained and performed, 80 of them were positive for the development of microorganisms, which represents 51.6%, on the other hand, 48.4% representing 75 cultures that did not present microbial development.

Of the 51.6% of positive cultures, 35 strains of *E coli* were found, equivalent to 46.66% of which 45.71% are *E coli* strains that produce Extended Spectrum Betalactamases. 20 strains of *E* faecalis representing 26.6%, 3 isolates of *S agalactiae* 4%, 2 isolates of *E cloacae, C. yungae* and *P milabiris* respectively equivalent to 2.66% for each strain. And 1 isolate of *P aeruginosa, K pneumoniae, G vaginalis, E aerogenes, C spp, C freundii, Aeromonas spp, S saprophyticus* and *S anginosus* that represent 1.33% each bacterial strain. In addition, there was development of 2 cultures by the yeast *C albicans* 2.66%

The strains show three types of sensitivity to quinolones; susceptible strains, strains with intermediate sensitivity and resistant strains shown in the table 1,

**Table 1.** Isolated Uropathogens and Their Resistance Patterns to Ciprofloxacin and Levofloxacin

| Table 1 | Ciprofloxacin |  |  | Levofloxacin |  |  |
| --- | --- | --- | --- | --- | --- | --- |
|  | Resistant | Intermediate sensitivity | Sensitive | Resistant | Intermediate sensitivity | Sensitive |
| *E coli* non BLEE | 5 | 0 | 14 | 5 | 2 | 12 |
| *E coli* BLEE | 13 | 0 | 3 | 14 | 2 | 0 |
| *E faecalis* | 2 | 0 | 18 | 1 | 1 | 18 |
| *S agalactiae* | 1 | 1 | 1 | 0 | 0 | 3 |
| *E cloacae* | 0 | 0 | 2 | 0 | 0 | 2 |
| *C. yungae* | 2 | 0 | 0 | 2 | 0 | 0 |
| *P milabiris* | 2 | 0 | 0 | 2 | 0 | 0 |
| *P aeruginosa*, | 1 | 0 | 0 | 1 | 0 | 0 |
| *K pneumoniae*, | 1 | 0 | 0 | 0 | 1 | 0 |
| *G vaginalis* | 1 | 0 | 0 | 1 | 0 | 0 |
| *E aerogenes* | 0 | 0 | 1 | 0 | 0 | 1 |
| *Citrobacter spp,* | 1 | 0 | 0 | 1 | 0 | 0 |
| *C freundii* | 1 | 0 | 0 | 1 | 0 | 0 |
| *Aeromonas spp,* | 0 | 0 | 1 | 0 | 0 | 1 |
| *S saprophyticus* | 0 | 0 | 1 | 0 | 0 | 1 |
| *S anginosus* | 1 | 0 | 0 | 0 | 0 | 1 |

The results were divided to evaluate which of the two antibiotics studied had greater resistance. For ciprofloxacin there are 27 resistant strains 37%, 1 strain with intermediate resistance 1% and 45 susceptible strains 62%. For levofloxacin 26 strains are resistant 36%, 41 strains are sensitive 56% and 6 strains show intermediate sensitivity 8%. Graph 1 and Graph2

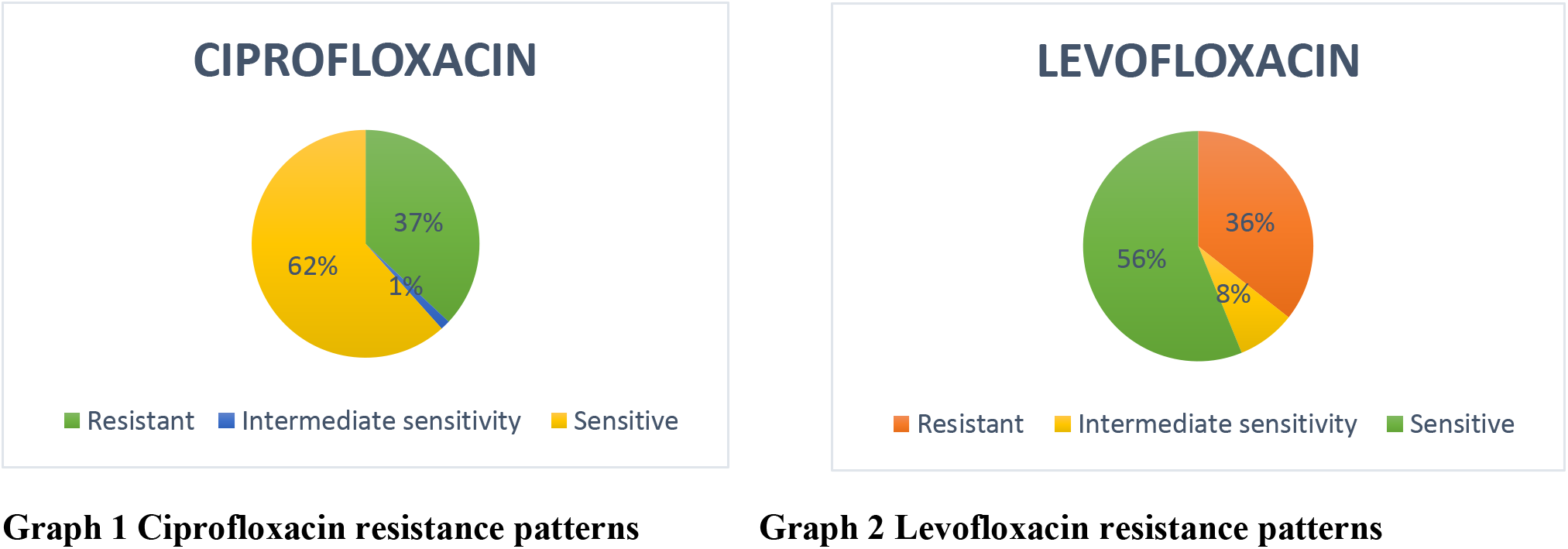

It was also identified in which bacteria had greater resistance, if in Gram positive or Gram negative results, obtaining the following results: For ciprofloxacin 4 Gram positive strains showed resistance, 20 strains are sensitive and 1 strain showed intermediate resistance. For the same antibiotics but in gram-negative bacteria, there are 23 resistant strains and 23 sensitive strains. Graph 3

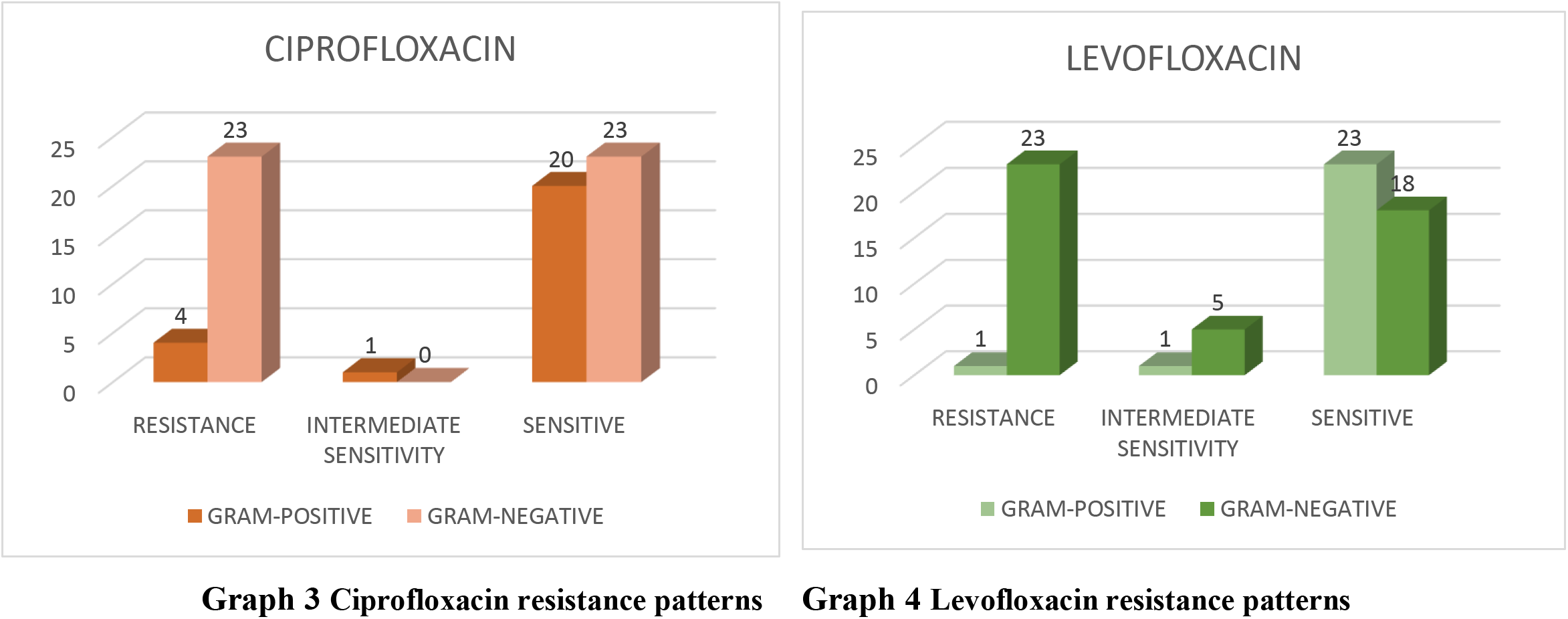

For levofloxacin, in gram-positive bacteria there are 1 resistant strain, 23 susceptible strains and 1 intermediate susceptible strain. For the same antibiotic but now with gram-negative bacteria, there are 23 resistant strains, 18 susceptible strains and 5 strains with intermediate sensitivity. Graph 4

## Discussion

The indiscriminate and extensive use of quinolones both in animal production and care, as well as in human medicine is associated with a high rate of resistance by bacteria to these antibiotics in an emergent and accelerated manner.

As can be seen in the following graphs, gram-negative bacteria are those with the highest resistance to ciprofloxacin and levofloxacin with resistant strains 23 respectively, while for gram-positive bacteria only 23 and 18 strain respectively they present resistance to ciprofloxacin and levofloxacin respectively. Giving a total of resistant strains between gram positive and gram negative of 30 to cipofloxacin and 24 to levofloxacin

Quinolone resistance is caused by various mechanisms, particularly plasmid-ediated quinolone resistance (PMQR) which contains the pentapeptide repeat family Qnr proteins (QnrA, QnrB, QnrS, QnrC, and QnrD). These proteins confer quinolone resistance by physically protecting DNA gyrase and topoisomerase IV from quinolone acts. This condition may provide a selective advantage for the development of quinolone resistance which could result in therapeutic failure. 1

In the present research, the bacterium that was mostly isolated and is the bacterium that occupies the first place in urinary tract infections was *Escherichia coli* with 35 strains, of which 19 are non-ESBL producers and 16 strains are ESBL producers.

In Escherichia coli, resistance to quinolones frequently occurs through mutation in gyrA and less often by gyrB genes, which catalyzes ATP dependent DNA supercoiling. Some other mechanisms of *E. coli* resistance to quinolones and fluoroquinolones are through efflux pumps and reduced drug accumulation in the bacteria due to changes in the purine protein. Many studies have revealed that mutations in a small parts of gyrA N-terminal (Amino acids 67 (Ala-67) to 106 (Gln-106)) leads to quinolones and fluoroquinolones resistance which is named quinolone resistance-determining region (QRDR).2

Producing β-lactamase enzymes are the most common mechanism of bacterial resistance. Extended-spectrum β-lactamases (ESBLs)-producing bacteria have usually multi-drug resistance, because most of the times, the genes related to the other resistive mechanisms have been also placed on the same plasmid carrying the genes encoding ESBLs. 3

In the present study, the *Escherichia coli* strains that produce extended spectrum beta-lactamases show a high pattern of resistance to both ciprofloxacin and levofloxacin with 13 and 14 strains respectively, unlike *Escherichia coli* that does not produce extended spectrum beta-lactamases where only 5 strains are resistant to both antibiotics.

For the rest of the isolated bacteria, the resistance patterns are variable, showing a greater tendency to sensitivity to levofloxacin and ciprofloxacin.

## Conclusion

155 urine cultures were performed in patients with suspected urinary tract infections, of which 80 were positive for bacterial growth, the objective being the evaluation of levfloxacin and ciprofloxacin on isolated bacteria, showed a high tendency in gram-negative, and even higher in extended spectrum beta-lactamase producing strains. On the other hand, gram positive strains tend to show different resistance patterns, with a greater tendency to sensitivity to both antibiotics. However, the rapid increase in resistance to broad spectrum antibiotics such as those evaluated in this study is alarming.

## Ethical approval

The following study was approved by the ethical committee of the Microtec laboratory

No permission of the patients was required for the obtention of the clinical isolates since they are no human material and are routinely recovered from patients cultures.

## Interest Conflict

The authors declare that they have no conflict of interest.

## Financing

No funding was received to carry out this work.

## Acknowledgment

We thank all the institutions that allowed us to carry out this research work. And dedicate this work to Victoria daughter

## Notes

### Competing Interest Statement

The authors have declared no competing interest.

